# Machine learning-based meta-analysis of colorectal cancer and inflammatory bowel disease

**DOI:** 10.1101/2023.08.04.551970

**Authors:** Aria Sardari, Hamid Usefi

## Abstract

Colorectal cancer (CRC) is a major global health concern, resulting in numerous cancer-related deaths. CRC detection, treatment, and prevention can be improved by identifying genes and biomarkers. Despite extensive research, the underlying mechanisms of CRC remain elusive, and previously identified biomarkers have not yielded satisfactory insights. This shortfall may be attributed to the predominance of univariate analysis methods, which overlook potential combinations of variants and genes contributing to disease development. Here, we address this knowledge gap by presenting a novel multivariate machine-learning strategy to pinpoint genes associated with CRC. Additionally, we applied our analysis pipeline to Inflammatory Bowel Disease (IBD), as IBD patients face substantial CRC risk. The importance of the identified genes was substantiated by rigorous validation across numerous independent datasets. Several of the discovered genes have been previously linked to CRC, while others represent novel findings warranting further investigation.

## Introduction

Colorectal cancer (CRC) ranks as one of the top three deadliest cancers worldwide, with an estimated 1.8 million cases and 881,000 fatalities in 2018 alone [1]. Timely detection of CRC can significantly improve prognosis and reduce mortality rates [2]. When CRC is diagnosed in individuals below the age of 50, it is referred to as early-onset CRC (eoCRC). Over the past few decades, the epidemiology of eoCRC has been subject to change, as reported by numerous studies. Starting from the 1990s, there has been a rise in the incidence of eoCRC across the world, including both high- and low-income countries [3, 4]. The rate of increase in eoCRC incidence is accelerating and is predicted to pose a significant public health challenge [3, 4]. Recently, the US Preventive Services Task Force recommended lowering the average-risk population screening age to 45 years [5, 6]. Possible justifications for the increasing incidence of eoCRC include a westernized diet, including red and processed meats; consumption of monosodium glutamate, titanium dioxide, high-fructose corn syrup and synthetic dyes; obesity; stress; and widespread use of antibiotics [7].

Due to its heterogeneity, CRC is controlled by many genes and environmental factors. Epigenetics refers to alterations in gene expression or function without changes in DNA sequence. Primary epigenetic modifications include DNA methylation, post-transcriptional modifications of histone and non-coding RNA-mediated changes of gene expression [8]. Despite its significant recognition, the contribution of epigenetic events to cancer evolution needs further investigation [9, 10]. It is believed that the modifications in epigenetics and the changes in the expression of non-coding RNAs can be utilized as biomarkers for the diagnosis, prediction of treatment response and prognostication in the case of CRC [11]. The genetic and epigenetic modification of cancer-associated genes occurs independently but recurrently in CRCs, and that epigenome alterations probably control important tumour cell phenotypes, including escape from immune surveillance [12].

Recent studies have provided important insights into the molecular mechanisms that underlie the formation of CRC. The majority of CRC cases (75%) are sporadic, while the remaining cases are either linked to inflammatory bowel diseases (IBD) or have a familial origin [13]. It is estimated that the process of CRC tumorigenesis is slow, taking almost two decades for a tumor to form [14]. Despite extensive research efforts and the elucidation of some pathways and genes, a considerable unknown portion of these diseases persists. In particular, the dynamics and complex process of cancer cell invasion and metastasis is poorly understood [15–17].

Oncogenic transformation in CRC is known to be caused by the driver genes APC, KRAS, SMAD4, and TP53, which modulate global translational capacity in intestinal epithelial cells [18]. Given our present understanding of the intricate nature of cancer genomes, how cancer cells evolve over time under treatment, and how inhibiting targets affects the body, it is now advisable to move away from the one gene, one drug approach and embrace a ‘multi-gene, multi-drug’ model for making informed decisions regarding therapy [19]. In other words, the unidentified aspect of the disease may stem from the cumulative effects of multiple low-penetrance genes, which together pose a substantial risk [20]. To that end, there have been considerable interest in molecular subtype classification of CRC using gene expression data, hierarchical clustering, and machine learning [19, 21–23].

Machine learning (ML) techniques have demonstrated their efficacy in addressing biological queries [24, 25]. Owing to their notable accomplishments, the application of ML methods to biological data is expanding, revealing their considerable potential in tackling genetic problems such as the imputation of missing SNPs and DNA methylation states, disease diagnosis [24, 26], antibody development [27], and numerous other areas. The use of ML has demonstrated great potential in enhancing our comprehension of cancer dynamics, and it holds the possibility of substantially transforming our understanding of cancer dynamics by revealing fresh insights into the molecular mechanisms that drive cancer progression and impact treatment response [28, 29].

In this paper, our primary goal is to investigate the genetic landscape that underlies the progression of both CRC and IBD, with a specific emphasis on additive gene interactions. Figure 1 provides a schematic representation of the research pipeline and the various tasks executed for both IBD and CRC.

**Fig 1.**
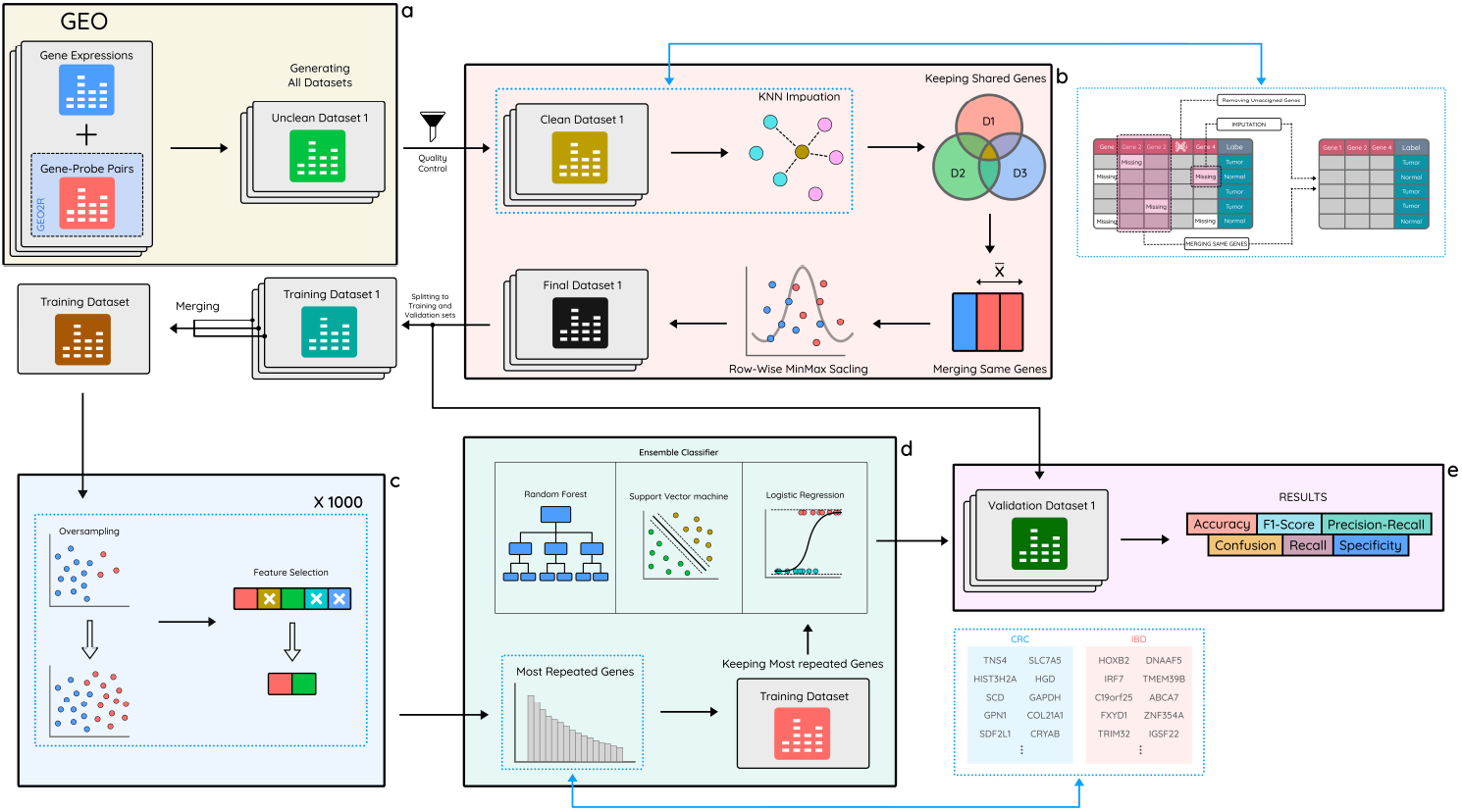
Schematic representation of the research workflow. **a** Raw datasets are retrieved from GEO, and tabular datasets are generated utilizing gene expression data and probe-gene mapping. **b** Data processing steps are performed, including discarding unassigned genes, imputing missing values, removing non-common genes, combining identical genes, and scaling each dataset. **c** After splitting datasets into training and validation sets and merging training sets to form a single set, a 1000-iteration oversampling/feature selection process is applied to identify the most prominent genes. **d** An ensemble classifier, comprising Random Forest, Support Vector Machine, and Logistic Regression, is trained on the training set. **e** The results are validated on the validation sets using the trained model, and four performance metrics – accuracy, F1-score, precision-recall, and confusion matrix – are employed for the evaluation of case-control sets and specificity and recall are employed for case-only and control-only sets.

We employed novel ML algorithms trained on case-control datasets from the GEO (Gene Expression Omnibus) database, consisting of 566 CRC cases and 262 controls. Through this process, we identified a subset of prominent genes capable of cumulatively distinguishing between CRC and control samples. To demonstrate the efficacy of our selected genes, we conducted validation using the top 40 genes on multiple external and independent datasets. Remarkably, our model accurately classified 1807 out of 1860 cases and 169 out of 173 controls, highlighting the strength of our approach. Although some of the chosen top genes were already known and studied in the literature, it is noteworthy that building a model solely based on just those well-known genes did not yield satisfactory validation results, leading to numerous misclassification.

Additionally, recognizing the heightened risk of CRC development in IBD patients, we also set out to identify a subset of genes capable of distinguishing between IBD cases and healthy controls. To accomplish this, we trained ML algorithms on GEO datasets comprising 288 IBD cases and 76 controls. Using the top 100 selected genes, our validation on external IBD datasets led to the correct classification of 212 out of 231 IBD cases and 51 out of 54 healthy controls. We note that the misclassified samples included 9 inflamed IBD samples that were misclassified as healthy.

With the aim of bridging the gap and identifying intermediary genes between CRC and IBD, we leveraged the STRING (Search Tool for the Retrieval of Interacting Genes/Proteins) platform to construct a gene network, integrating the top CRC and IBD genes. Intriguingly, we observed direct interactions between several IBD-associated genes and CRC-associated genes without the need for intermediary genes. Notably, the GAPDH gene emerged as a pivotal player in linking the two gene subsets and proved to be one of the most crucial CRC-associated genes identified in our study.

Our findings warrant further investigation into the role of the oncogene TNS4, as well as SLC7A5 and SCD, in the nuclear receptors meta-pathway. Additionally, our other top selected genes, namely SLC7A5, SCD, GAPDH, and SDF2L1, are involved in the mTOR signaling pathway and merit thorough exploration. These valuable insights significantly contribute to enhancing our understanding of the intricate genetic mechanisms underlying the progression of CRC and IBD.

## Results

### Data retrieval and curation

We combined 6 gene expression datasets from the GEO (Gene Expression Omnibus) database to form a training dataset, and an additional 14 different gene expression datasets were selected for validation. Some of the validation datasets contain only cases. A comprehensive summary of each CRC dataset is presented in Supplementary Table 1 for further reference. All datasets in this paper consist of biopsy samples, and no blood samples are included. We carefully examined all the validation datasets to make sure there is no overlaps or leakage with training datasets. For instance, dataset GSE32323 was omitted due to a high probability of containing identical patients (with differing expressions) as those in dataset GSE21510. Additionally, to enhance reliability, gene expression samples were grouped based on geographical similarity (country/city) within either the training or validation sets as much as possible. Notably, GSE44076 contains samples from healthy donors in addition to tumor samples and matched normal samples. Regarding this dataset, we also used healthy donors to see if our model could diagnose healthy samples because it was trained on tumour and non-tumour samples. It is important to note that datasets GSE68468, GSE103512 and GSE2109 encompass samples derived from various organs in addition to the colon and rectum; however, in our analysis exclusively, we only included samples derived from colon and rectum. To be able to merge the training datasets together and then perform the validation, we only kept genes that are common between all training and validation datasets. In the end, all the CRC datasets had uniformly 10,113 genes.

We selected 5 gene expression IBD datasets from GEO for training and 6 different gene expression IBD datasets for validation. We included only those samples who had not undergone any specific treatment. Additionally, we excluded datasets consisting of blood samples. Dataset GSE16879 includes pre- and post-infliximab treatment samples, of which we selected only the pre-treatment ones. In GSE59071 and GSE48958, inactive samples were excluded. From GSE179285, we included only inflamed samples and from GSE4183, only IBD and normal samples were chosen. From GSE37283, patients diagnosed with ulcerative colitis with neoplasia were retained. Supplementary Table 2 contains details of the datasets used for IBD. After discarding genes that are not common between all IBD datasets, all IBD datasets had uniformly 16,413 genes.

### ML models, evaluation, and performance metrics

One of the most pivotal elements of any ML model applied to genomic data is feature selection which is tasked with uncovering the most disease-relevant genes among thousands. The detection of low-penetrance genes related to CRC and IBD necessitates the use of wrapper or hybrid feature selection techniques. However, due to the computational complexity, wrapper methods are not feasible for high-dimensional datasets like those employed in this paper. In this research, we utilized SVFS (Singular Vectors Feature Selection), a hybrid feature selection method that has recently demonstrated superior results compared to other methods on gene expression data [30]. As we can see from In Supplementary Tables 1 & 2, the number of cases is much more than the number of controls. To avoid developing biased models, we employed SMOTE [31] (synthetic minority oversampling technique) to use the controls in the training datasets and generate synthetic controls; this way, we equalize the number of cases and controls within the training dataset. We run the SVFS on the training dataset to select the first 100 most important genes, as shown in Supplementary Tables 3 & 4.

To build an ML model, we employed an ensemble classifier consisting of Random Forest (RF), Logistic Regression (LR), and Support Vector Machine (SVM). Each RF, LR, and SVM are based on different algorithms, and their ensemble provides greater robustness than single models and decreases the potential for overfitting. The ensemble classifier was trained on the reduced training dataset (using only the most important genes) to build a model. Finally, the model was evaluated on all validation datasets. We employed accuracy, the precision-recall curve, and the confusion matrix to report validation results. Figure 2 illustrates the validation results for each CRC case-control validation dataset. We note that using only the first 40 genes for model generation, the model comes close to achieving optimal performance on all validation sets. Confusion matrices and precision-recall curves are generated based on these 40 genes.

**Fig 2.**
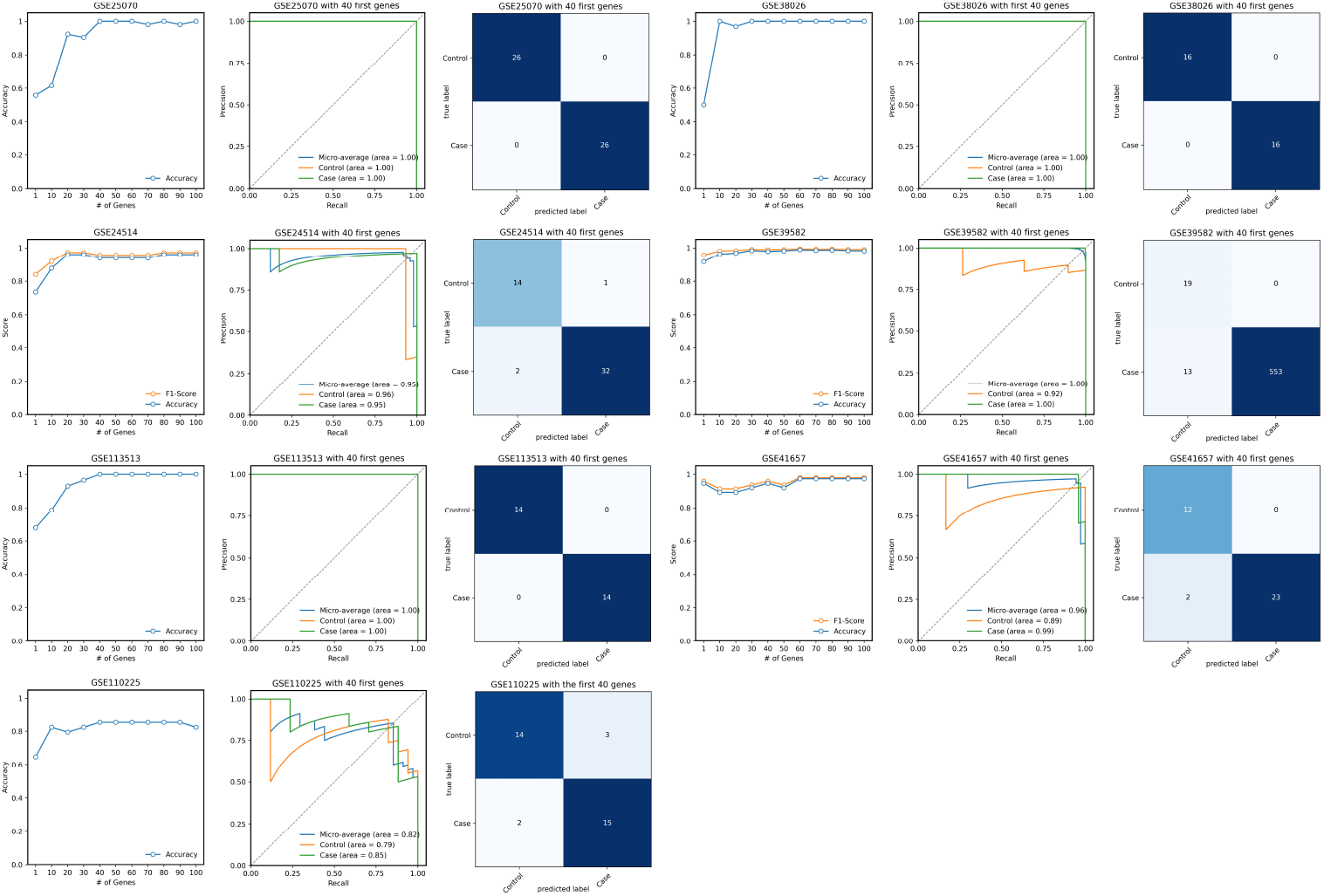
Evaluation of identified CRC genes on independent validation sets. Accuracy and F1-score are plotted for the different number of prominent genes utilized for training and validation. Confusion matrices and precision-recall curves (including AUC) are plotted using the first 40 prominent genes.

Several of our datasets consisted of cases only. So, we used the F1-score to report the validation results on these datasets. For case-only datasets, we adopted recall (also known as sensitivity or true positive rate), representing the proportion of correctly predicted case samples relative to the overall number of cases. We further incorporated 51 healthy donors from dataset GSE44076 as another validation set to evaluate the final model’s ability to identify healthy donors. For this dataset, we utilized specificity, which, in this case, is defined as the ratio of correctly predicted control samples to the total number of controls. Figure 3, illustrates the results for case-only datasets as well as results for healthy controls of the GSE44076 dataset.

**Fig 3.**
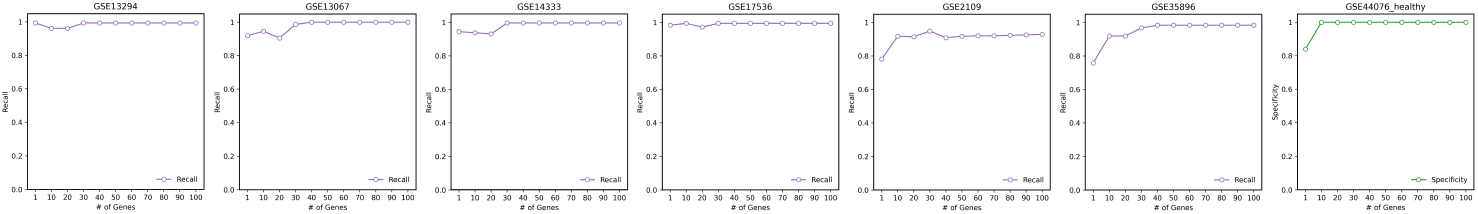
Evaluation of the model trained on tumor and matched normal samples on case-only and control-only datasets.

In order to achieve reliable results from ML algorithms, it should be noted that the validation datasets must not be utilized at any point during the training or model generation process. Given the validation results in Figures 2 and 3, we deduce that using 40 prominent identified genes, our model could diagnose 1807 cases out of 1860 and 169 controls out of 173 (including healthy samples from GSE44076). Some of the previously reported significant genes include TP53, APC, KRAS, MGMT [32], SMAD2 [33] and SMAD4 [33]. It is interesting to note that if we build a model just based on these well-known genes, we do not get acceptable validation results. Indeed, we implemented a supplementary pipeline using only TP53, APC, KRAS, MGMT, SMAD2, and SMAD4, and it turned out that 109 controls out of 153 are misclassified, which is a poor performance (for this experiment, we had to exclude GSE38026 because the KRAS gene does not exist in this dataset).

The same methodology was employed for IBD; that is, SVFS was utilized on the training IBD dataset, and the first 100 significant genes were selected, as shown in Supplementary Table 4. As demonstrated in Figure 4, the ensemble classifier effectively distinguished inflamed samples from healthy samples in GSE9452, GSE37283, GSE4183 and GSE48958. In the case of GSE36807, the model accurately diagnosed all healthy samples, though nine inflamed samples were misclassified as healthy. Overall, the classifier exhibited strong performance, suggesting an acceptable identification of IBD-related genes by identifying 212 IBD cases out of 231 and 51 healthy controls out of 54.

**Fig 4.**
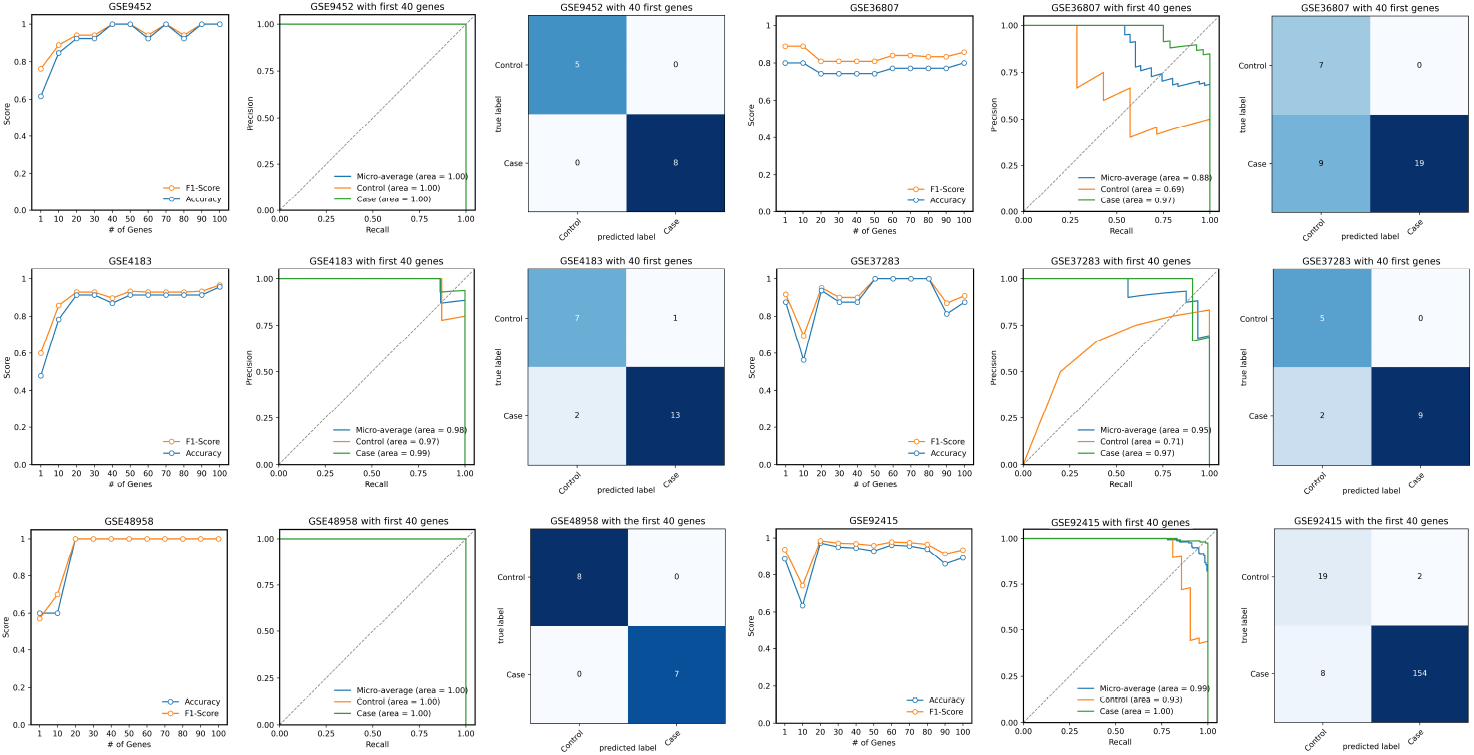
Evaluation of identified IBD genes on independent validation sets. Accuracy and F1-score are plotted for the different number of prominent genes utilized for training and validation. Confusion matrices and precision-recall curves (including AUC) are plotted using the first 40 prominent genes.

### Analyzing Gene Interactions

In order to discover and comprehend the inherent interactions among the identified genes, we utilized STRING (Search Tool for the Retrieval of Interacting Genes/Proteins). We constructed a gene network by integrating the top 50 IBD genes with 50 CRC genes in the initial step, setting the interaction score to medium confidence. Figure 5a illustrates the resulting network.

**Fig 5.**
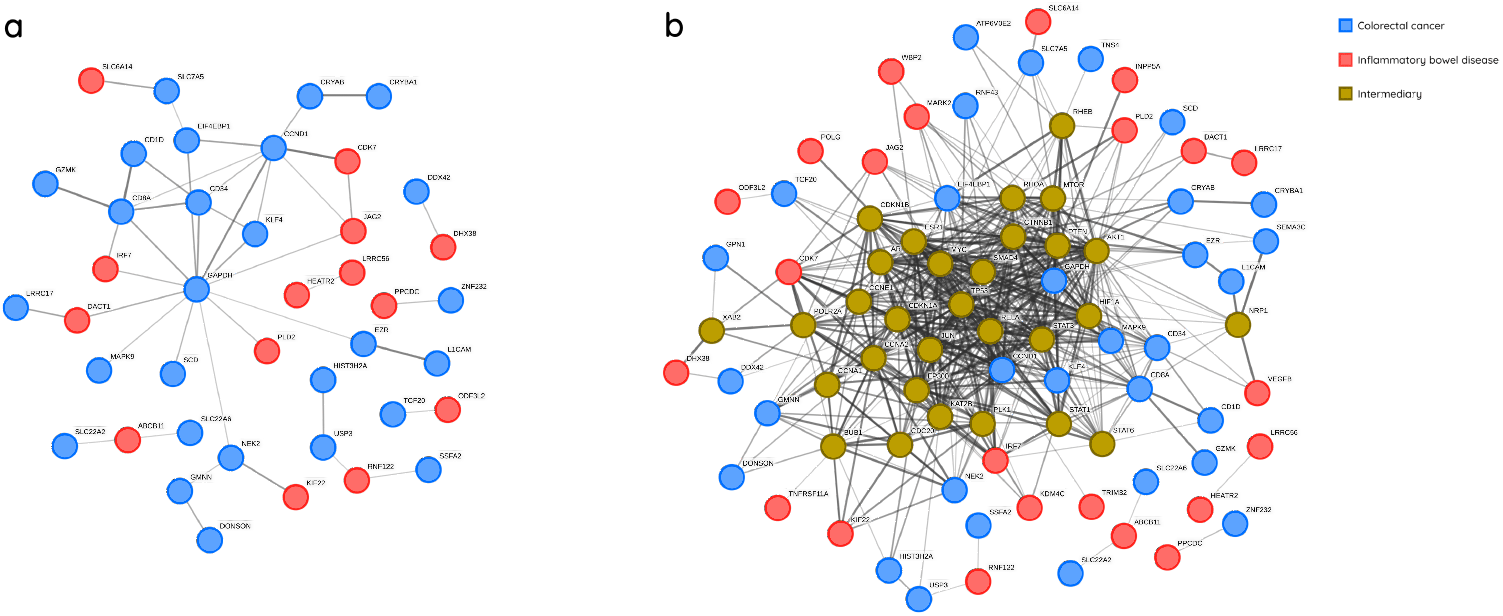
IBD and CRC gene interaction networks generated by STRING for identified genes. **a** Network generated based on CRC and IBD genes without the participation of intermediary genes. **b** Network generated based on CRC and IBD genes with the participation of intermediary genes.

As observed in Figure 5a, 13 IBD-associated genes and 27 CRC-associated genes have direct interaction (without intermediary genes). The GAPDH gene appears to play a pivotal role in linking the two gene subsets and is one of the most crucial CRC-associated genes identified in our subset. To investigate potential interactions between CRC and IBD genes, we extended the network in Figure 5a to include intermediary genes that may serve as a bridge between CRC and IBD genes. For this extended network in Figure 5b, we took into account only single intermediary genes, which is a drawback since the bridge could involve two genes, for example. While single intermediary genes are more influential, other genes with minor additive effects are overlooked. Supplementary Table 5 lists the full names of single intermediary genes. Owing to the network’s complexity, we preserved genes in Figure 5b with a higher number of connections in the network for illustrative purposes. For example, TP53 may be regarded as the most critical intermediary gene. This gene and its adjacent genes might be fundamental to IBD and CRC progression. The intermediary genes are highly likely to contribute to the disease due to their strong connections to genes in our subset and, importantly, their bridging functions. We further explored the COSMIC (Catalogue Of Somatic Mutations In Cancer) database to determine if any of the genes in our subset had been previously reported as having a strong association with CRC. Tissue selection, Sub-tissue selection, Histology selection, and Sub-histology selection were set to Large intestine, Include all, Carcinoma, and Adenocarcinoma, respectively. Remarkably, TP53, SMAD4, RNF43, CTNNB1, and PTEN ranked among the top 20 most frequently mutated CRC-related genes listed in COSMIC. On the other hand, several of our reported significant genes were not on the COSMIC list and did not receive adequate attention from researchers.

We also performed GSEA (Gene Set Enrichment Analysis) using most first 50 identified CRC genes. GSEA showed that TNS4, SLC7A5, and SCD are involved in the nuclear receptors meta-pathway. Genes SLC7A5, SCD, GAPDH and SDF2L1 were involved in the mTOR signalling pathway. Also, several other gene sets were associated with cell cycle regulation and transcription regulation. Given the limited understanding of the underlying mechanisms of CRC and IBD, we propose to consider other important genes that are not part of the network in Figure 5. For instance, an in-depth investigation of TNS4, GAPDH, L1CAM, GAL, CRYAB, IRF7, GPN1, TMEM39B, EZR, and all other genes referred to in Supplementary Tables 3 and 4 are needed to discern their role in CRC and IBD.

## Discussion

One of the customary approaches to find potential biomarkers for a disease is to identify genes that are differentially expressed (DEG) in cases and controls [34–37]. Such methods usually set a threshold to identify DEGs, and as such many genes are filtered out even if they were close to the threshold. Even though one can validate whether a candidate gene is DEG on an external dataset, usually, no single gene can be the sole cause of cancer, and there is no methodology to quantify the relation of a subset of DEGs to cancer. In contrast, machine learning and feature selection algorithms, when validated on external datasets, can identify promising biomarkers.

In this study, we identified several potential genes correlated with CRC using multiple multivariate machine-learning methods. Some of these genes have not received enough attention in previous studies. Our findings provide new insights into the potential additive genes correlated with CRC and IBD, as patients with IBD are at high risk of developing CRC. Identifying novel genes correlated with CRC and IBD provides a foundation for future research into the underlying mechanisms of these diseases.

Notably, TNS4 (CTEN) emerges as a critical gene associated with CRC. Despite previous studies highlighting TNS4’s association with CRC, its significant contribution has been relatively overlooked. This gene has been identified as an oncogenes gene in several studies. It has been argued that TNS4 plays a pivotal role in CRC tumorigenesis, suggesting that TNS4 suppression could represent a promising therapeutic approach [38] and its knockdown improves sensitivity to Gefitinib [39]. It has been revealed that TNS4 interacts with MET signalling [40], a process known to enhance motility, cancer cell survival, and angiogenesis [41]. This interaction between TNS4 and MET is direct, and TNS4 plays a role in maintaining MET stability, thereby supporting cancer cell survival [40].

We also identify GAPDH is another potential CRC-related protein-coding gene. Not only does it connect a significant number of CRC and IBD-related genes (Figure 5b), but it also ranks among the top 40 most prominent CRC-related genes given in Supplementary Table 3. Investigations have examined GAPDH’s interaction with mutated KRAS and BRAF, suggesting that GAPDH suppression via vitamin C may disrupt tumor growth [42]. Other work has analyzed tumor versus non-tumor pairs in 195 cases, identifying substantial overexpression of GAPDH in CRC cases [43]. Researchers have also observed significant upregulation of GAPDH in CRC, indicating its potential value in early CRC detection [44].

SLC7A5 is identified as the second most significant gene on our list (Supplementary Table 3). Najumudeen et al. conducted comprehensive research on SLC7A5’s correlation with CRC [45]. They proposed that SLC7A5 might offer potential therapy for KRAS-mutant CRC unresponsive to other treatments [45]. Additionally, Huang et al. identified SLC7A5 as one of the five key genes involved in the ferroptosis of colon cancer cells [46].

HIST3H2A, HGD, GPN1, COL21A1, and SDF2L1 appear as potential novel biomarkers using out results. While there is evidence linking HIST3H2A to lung cancer [47] and pancreatic cancer [48], its association with CRC remains unverified. Yi et al. found a significant association between HGD and rectal cancer [49], but no other research has established a strong link between HGD and CRC. To our knowledge, GPN1 is a novel gene identified through our work. Presently, limited information is available on this crucial gene, warranting further investigation for a more comprehensive understanding. As per our review, Li et al.’s study stands as the only research that has identified COL21A1 as a possible diagnostic marker [50]. Despite examining the link between the SDF2L1 gene and different cancer types, including Nasopharyngeal Carcinoma [51], its potential role as a CRC marker has not been acknowledged.

Cruz-Gil et al. identified SCD as a critical component of lipid metabolism in CRC [52]. The relationship between SCD and ACSL increases the risk of relapse in CRC patients [52]. Furthermore, Liao et al.’s research suggests a connection between overexpressed SCD-1 and advanced CRC [53].

CRYAB has been linked to various cancer types [54], and multiple studies have explored its association with CRC. Deng et al. characterized CRYAB as a tumor-suppressor gene and a potential diagnostic marker [55], and Shi et al. verified its function as a prognostic CRC biomarker [54]. Dai et al. also proposed CRYAB as a promising target for CRC therapies [56].

Another dimension of importance is the role of non-coding RNAs in CRC. It is believed that modifications in epigenetics and alterations in the expression of non-coding RNAs can be harnessed as biomarkers for CRC diagnosis, prognostication, and treatment response prediction [11]. Non-coding RNAs, especially microRNAs and long non-coding RNAs, play significant roles in gene expression regulation and are implicated in several CRC pathways [57–59]. For instance, RAMS11, a non-coding RNA, is highlighted for its regulation of topoisomerase II*α* (TOP2*α*), underscoring its potential value as a biomarker and therapeutic target for metastatic CRC [60]. The increased expression of lncRNA IGFL2-AS1 in CRC tumor tissues and cells [61] further supports this perspective. By concentrating predominantly on coding regions of DNA, we might have overlooked these crucial actors in CRC’s genetic landscape. Incorporating both coding and non-coding DNA segments could provide a more comprehensive insight into CRC’s genetic intricacies, thereby shaping its diagnosis and treatment avenues more effectively.

Validation results on CRC and IBD datasets provided us with a set of genes that are deemed important in the development of CRC and the transition of IBD into CRC. Our findings offer new insights into the potential additive genes correlated with CRC and IBD and emphasize the value of machine learning algorithms in identifying genes that may contribute to CRC and IBD development. Nonetheless, our work here has certain limitations, such as the utilization of individual intermediary genes to establish connections between significant genes associated with IBD and CRC. We also note that many genes were removed at the initial stage of our preprocessing procedure to obtain an identical subset of genes across all datasets.

Further studies focusing on genes’ additive functions instead of single-variate analyses are necessary to confirm these genes’ contributions. The potential significance of these findings for clinical applications includes the possibility of developing better prevention, detection, and treatment methods for CRC and IBD patients.

## Methods

### Data preprocessing

The presence of missing values in data can disrupt numerous machine learning algorithms’ functionality, potentially leading to biased outcomes, inaccurate predictions, and diminished accuracy. Addressing incomplete data is thus vital before deploying machine learning algorithms, including those employed in the current study, through either the removal of incomplete observations or the imputation of missing values. The datasets used in this research contain several missing values. Discarding columns with missing values might result in losing vital disease-associated genes, and eliminating samples with missing values could compromise feature selection and classifier model performance due to reduced sample size. To address these missing values, we employed the K-Nearest Neighbors (KNN) imputation algorithm, with the number of neighbors set to five.

Numerous probes were assigned to the same genes after matching probe IDs with corresponding genes from the GEO2R mapping file. To integrate these probes into a single gene, the mean gene expression values were utilized. In order to integrate training datasets and execute uniform validation on validation sets, we maintained common genes across all datasets. Consequently, 10,113 and 16,413 genes were retained for CRC and IBD analyses, respectively.

Since we are merging several datasets from different populations and platforms, we need to scale the samples to ensure uniformity. In order to identify the optimal scaler, a variety of scalers were applied to the data. Principal component analysis (PCA) was utilized to reduce data dimensionality and visualize the outcomes for each scaler. Figure 6 demonstrates that both row-wise MinMax normalization and quantile normalization yield improved dispersion of CRC datasets. Consequently, the model is better equipped to discern the underlying data patterns related to gene contributions. Given the varying ranges of different genome datasets, row-wise MinMax normalization offers an advantage over a quantile transformer when classifying new datasets or samples. Furthermore, it preserves gene expression correlations while transforming gene expression into a range of 0 to 1 for each instance. Hence, row-wise MinMax scaling was executed prior to feature selection across all datasets. Row-wise MinMax scaling was also applied to transform the IBD datasets.

**Fig 6.**
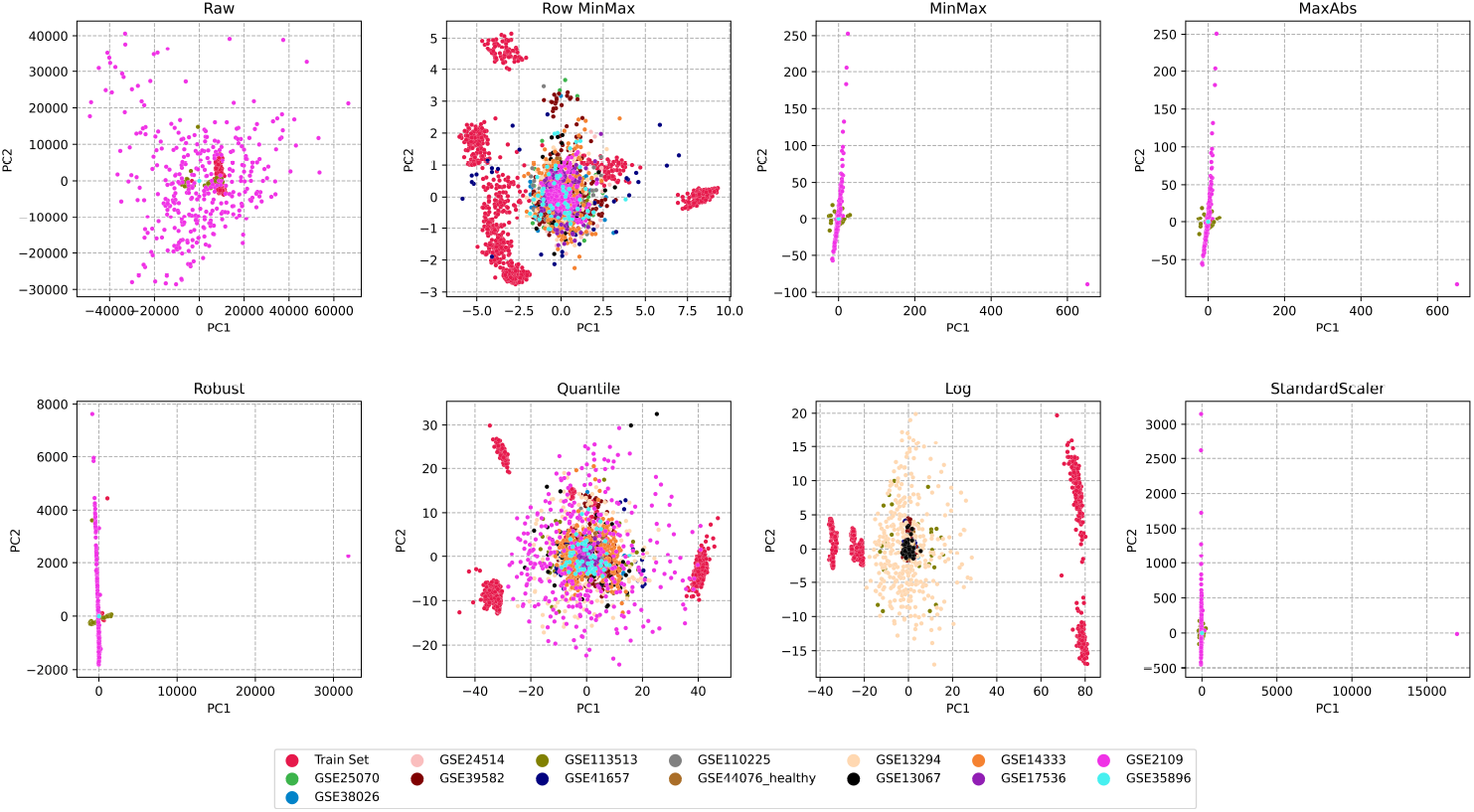
Effect of different scalers on CRC training dataset and test datasets.

We also note that the number of cases is much more than the number of controls in both our CRC and IBD training datasets. The high case-to-control ratio undermines the performance of both feature selection and the classification model and can build models that are biased toward cases. To tackle this issue, we employed SMOTE [31] (synthetic minority oversampling technique), an effective oversampling strategy, to use the controls in the training datasets and generate synthetic controls; this way, we equalize the number of cases and controls within the training dataset.

### Feature selection

We incorporated the parameters suggested for biological data in the paper as parameters for our algorithm (Th_irr_ = 3, Th_red_ = 4, *α* = 50, *β*=5, and *k* = 100) [30]. Executing the SVFS feature selection algorithm may yield varying subsets of features on each iteration. To achieve consistent results and verify the relationship between the identified genes and the disease, we ran the algorithm 1000 times on the training dataset and identified the top 100 genes that were repeated the most throughout the 1000 runs. Oversampling impacts dataset structure and may alter the gene subset selection by the feature selection algorithm. Consequently, oversampling was conducted before each iteration of feature selection to guarantee a more robust selection. Supplementary Figure 1 illustrates the top 100 most repeated genes and their number of occurrences in the CRC (and IBD) train dataset. Once we determined the top 100 genes using SVFS on the training dataset, we discarded all other genes from all the datasets, including the training and validation. Then we trained our ensemble classifier (RF+LR+SVM) on the reduced train dataset. Finally, we validated the model on each of the validation datasets and reported several performance metrics. Among all the performance metrics, the confusion matrix is perhaps the most transparent one and all other metrics are derived from the confusion matrix.

**Table 1.** Gene Set Enrichment Analysis (GSEA) of top 50 CRC-related genes.

| Term | ES | NES | NOM *p*-value | FDR *q*-value | FWER *p*-value | Lead Genes |
| --- | --- | --- | --- | --- | --- | --- |
| mTORC1 signalling | 0.896 | 1.733 | 0.001 | 0.006 | 0.005 | SLC7A5, SCD, GAPDH, SDF2L1 |
| Cell cycle | -0.605 | -1.809 | 0.01 | 0.009 | 0.009 | FAP, USP3, DONSON<br>CCND1, KLF4, GMNN<br>EIF4EBP1, EZR, NEK2 |
| Nuclear receptors pathway | 0.868 | 1.706 | 0.001 | 0.0111 | 0.012 | TNS4, SLC7A5, SCD |
| Transcription regulator activity | -0.681 | -1.623 | 0.040 | 0.027 | 0.031 | ZNF232, CCND1, KLF4<br>ZXDC, GMNN |
| Cell cycle regulation | -0.578 | -1.627 | 0.028 | 0.092 | 0.046 | FAP, DONSON, CCND1<br>KLF4, GMNN, EIF4EBP1<br>NEK2 |

## Acknowledgments

This research was partially supported by grants from the Natural Sciences and Engineering Research Council of Canada (NSERC) to H.U. (grant number RGPIN: 2019-05650).

## Author contributions statement

Conceptualization H.U.; Methodology A.S. and H.U.; Coding A.S.; Experiments A.S. and H.U.; Writing A.S. and H.U.; Supervision H.U.

## Declaration of generative AI in scientific writing

During the preparation of this work the author(s) used Grammarly and ChatGPT in order to check the grammar and improve clarity. After using this tool/service, the author(s) reviewed and edited the content as needed and take(s) full responsibility for the content of the publication.

## Additional information

### Competing interests

The author(s) declare no competing interests.

### Use of experimental animals and human participants

This research did not involve human participants or experimental animals.

### Informed consent

Not applicable.

### Ethics approval

Not applicable.

## Notes

### Competing Interest Statement

The authors have declared no competing interest.

